# Assessing temperature-dependent competition between two invasive mosquito species

**DOI:** 10.1101/2020.06.24.167460

**Authors:** Michelle V Evans, John M Drake, Lindsey Jones, Courtney C Murdock

## Abstract

1. Invasive mosquitoes are expanding their ranges into new geographic areas and interacting with resident mosquito species. Understanding how novel interactions can affect mosquito population dynamics is necessary to predict transmission risk at invasion fronts. Mosquito life-history traits are extremely sensitive to temperature and this can lead to temperature-dependent competition between competing invasive mosquito species.
2. We explored temperature-dependent competition between *Aedes aegypti* and *Anopheles stephensi*, two invasive mosquito species whose distributions overlap in India, the Middle East, and North Africa. We followed mosquito cohorts raised at different intraspecific and interspecific densities across five temperatures (16°C - 32°C) to measure traits relevant for population growth and to estimate species' per capita growth rates. We then used these growth rates to derive each species competitive ability at each temperature.
3. We find strong evidence for asymmetric competition at all temperatures, with *Ae. aegypti* emerging as the dominant competitor. This was primarily due to differences in larval survival and development times across all temperatures that resulted in a higher estimated intrinsic growth rate and competitive tolerance estimate for *Ae. aegypti* compared to *An. stephensi*.
4. *Synthesis and applications:* The spread of *An. stephensi* into the African continent could lead to urban transmission of malaria, an otherwise rural disease, increasing the human population at risk and complicating malaria elimination efforts. Competition has resulted in habitat segregation of other invasive mosquito species, and our results suggest that it may play a role in determining the distribution of *An. stephensi* across its invasive range.

## INTRODUCTION

Multiple mosquito species have invaded new regions in recent decades (e.g, *Aedes albopictus* in the US and Europe (Medlock et al., 2012), *Culex coronator* in the US (Wilke et al., 2020), *Aedes japonicus* in Europe (Schaffner et al., 2009)). Invasive mosquito species often compete with native mosquito species for resources and have been implicated in the decline of native species in several instances (Kaufman and Fonseca, 2014; Lounibos et al., 2016). However, some invasive mosquito species are able to coexist with the native species in a portion of their range due to differences in competitive outcomes dependent on the environmental context (Lounibos and Juliano, 2018).

Context-dependent competition results when competition dynamics depend on the environmental context, particularly abiotic variables such as temperature (Chamberlain et al., 2014). For example, temperature-dependent competition has been observed in many systems and especially for temperature-sensitive organisms such as *Tribolium* beetles (Park, 1954), *Daphnia spp.* (Fey and Cottingham, 2011) and aphids (Grainger et al., 2018). Like most ectotherms, mosquito life history traits are highly dependent on temperature, and whether competition results in co-existence or exclusion may depend on temperature and species’ individual thermal niches.

Environmentally-dependent competition has affected the invasion dynamics of two mosquito species in the Southeastern United States, *Aedes aegypti* and *Ae. albopictus*. *Ae. aegypti* was introduced to the Americas in the 1700s (Brady and Hay, 2020), while *Ae. albopictus* was introduced more recently in the 1980s (Sprenger and Wuithiranyagool, 1986). Since its introduction, *Ae. albopictus* has reduced the range of *Ae. aegypti* in the southeastern US through a combination of female satyrization and larval competition (Lounibos and Juliano, 2018). However, larval competition is temperature-dependent, with *Ae. aegypti* able to persist at higher temperatures (Lounibos, 2002). The environmental-dependence of species coexistence is further complicated by the higher desiccation tolerance of *Ae. aegypti* eggs compared to *Ae. albopictus* (Juliano et al., 2002). In this case, temperature-dependent competition between *Ae. aegypti* and *Ae. albopictus*, in addition to asymmetric reproductive interference, led to habitat segregation and reduced population abundances of *Ae. aegypti* locally (Lounibos and Juliano, 2018).

Similar to the invasion of *Ae. albopictus* in the southeastern US, the range of *Anopheles stephensi* is currently expanding from its previous distribution within the Indian sub-continent and the Arab peninsula to parts of Northeast Africa, where *Ae. aegypti is* endemic (Seyfarth et al., 2019; Surendran et al., 2019). Unlike other *Anopheles* species that only breed in natural water bodies, *An. stephensi* breeds in artificial containers in urban areas (Thomas et al., 2016), where it serves as a primary vector of malaria (Singh et al., 2017). In these habitats, *An. stephensi* is often found co-habiting with other container species, including *Ae. aegypti* (Mariappan et al., 2015). Whether these species interact has not been studied. Both species demonstrate sensitivity to temperature, but their specific life history traits differ in their responses to temperature (Mordecai et al., 2017, 2013), which may translate to temperature-dependent differences in competitive ability. Additionally, *Ae. aegypti* is endemic to the African continent and a primary vector for arboviruses such as yellow fever and dengue, while *An. stephensi* is a primary vector of urban malaria in its endemic range. Understanding how these species interact can aid in predicting how the local mosquito vector community, and therefore human disease risk, may be affected by the invasion of *An. stephensi.*

We tested for the existence and measured the strength of temperature-dependent competition between *Ae. aegypti* and *An. stephensi* across a range of temperatures (16°C - 32°C). At each temperature, we followed cohorts of mosquitoes reared at different intra- and interspecific densities and measured life history traits relevant to population dynamics. We calculated per capita growth rates from these trait measurements and fit competition models to these data. From these models, we then calculated competition coefficients to estimate the relative competitive ability of each species at each temperature.

## Materials and Methods

### Experiment

We used a response surface design across fifteen density treatments (Fig. 1, Table S1) and five temperature treatments (16°C, 20°C, 24°C, 28°C, 32°C) to explore pairwise competition between *Ae. aegypti* and *An. stephensi*. Response surface designs vary the densities of competing species independently across a range of total densities, which allows for the fitting and parameterization of competition models (Inouye, 2001). Densities were chosen so that the median total density corresponded to standard rearing conditions for these strains (200 larvae/1 L of water), allowing for species densities that were lower and higher than optimum. The range of temperatures represented the full range at which each species can persist when reared individually, including field-relevant temperatures. The strain of *Ae. aegypti* was an outbred field-derived population originating from Tapachula, Chiapas, Mexico, 2016. The F5 generation was used in this experiment. The strain of *An. stephensi* (Liston) was sourced from a long-standing colony housed at Pennsylvania State University that was originally obtained from the Walter Reed Army Institute of Research. This experimental design was replicated three times.

**Figure 1.**
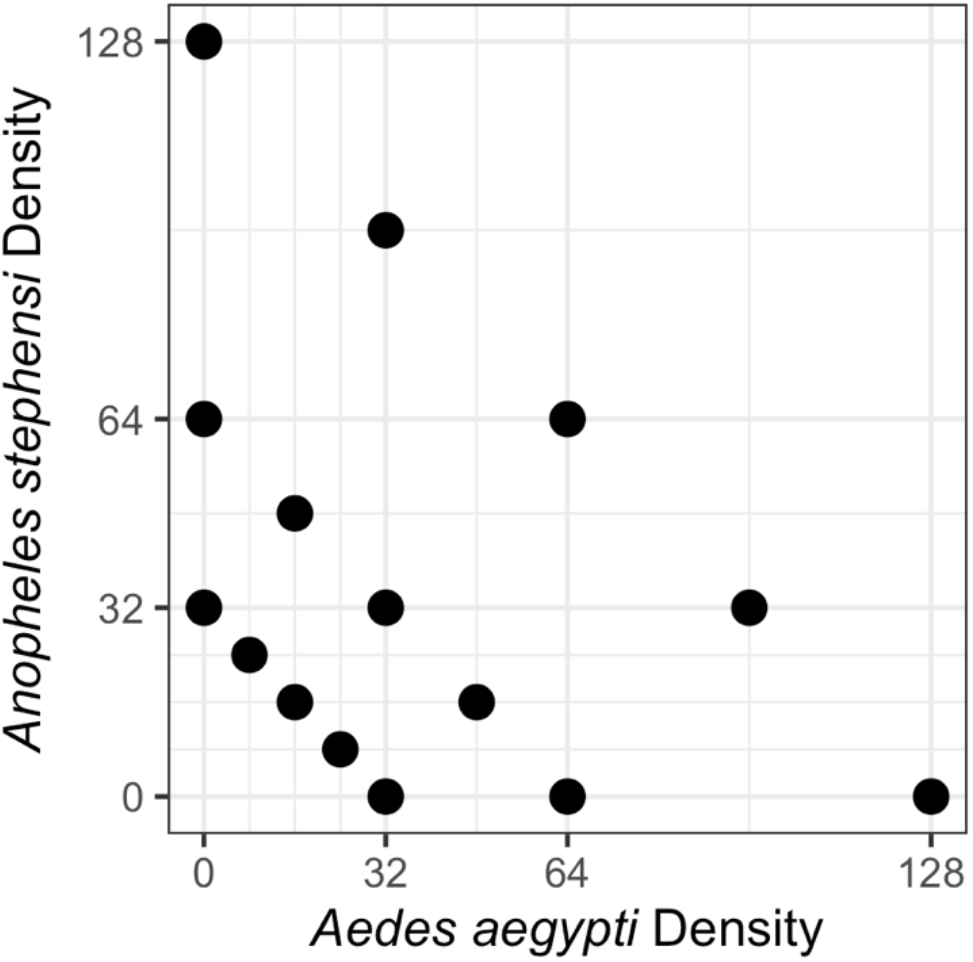
Species densities (per 250 mL) used in response surface design. Species densities are also noted in Table S1.

Larvae were hatched on experimental Day 0. On Day 1, 1^st^ instar mosquito larvae were placed in quart-size mason jars with 250mL of reverse osmosis filtered water and 0.1g cichlid pellet food (Hikari Cichlid Cod Fish pellets). Rearing jars were placed in incubators (Percival Scientific), following the intended temperature treatments with 85% (± 5%) relative humidity (RH), and 12:12hr light:dark diurnal cycle. Temperature regimens were programmed to a mean given by the experimental treatment (16°C, 20°C, 24°C, 28°C, 32°C ± 0.5°C) and daily periodic fluctuation of 9°C, following the Parton-Logan equation (Parton and Logan, 1981), which is characterized by a sine wave during the daytime and exponential curve at night. Rearing jars were inspected daily for emerged mosquitoes and the numbers of males and females emerging on each day recorded. Following emergence, adults were pooled by day of emergence, temperature, species, and density treatment. Adults were kept in a 16 oz paper cup in a walk-in incubator (Percival Scientific) at 27°C (± 0.5°C), 85% RH (± 5%), and 12:12hr light:dark cycle and offered a 10% sucrose solution *ad libitum*.

Mosquitoes that emerged up to and including the day of peak emergence were allowed to mate four to six days before being offered a blood meal. Forty-eight hours prior to blood feeding, the sucrose was removed and replaced with deionized water, which was then removed 24hr later. Blood meals were administered via a water-jacketed membrane feeder at 38°C for 30 minutes. A maximum of 10 blood-fed females per treatment were sorted into individual oviposition containers and kept at 27°C (± 0.5°C), 85% (± 5%) RH, 12:12hr dark:light cycle. Oviposition containers consisted of a 50mL centrifuge tube with a damp cotton ball and filter paper at the bottom to collect eggs. Centrifuge tubes were covered with a fine mesh to keep mosquitoes inside and allow for air circulation. During this time, females had access to a 10% sucrose solution *ad libitum*. Females were monitored daily for oviposition events. The date of the oviposition event was noted, and the number of eggs was counted the following day to allow for females who were monitored while ovipositing to finish laying eggs. After oviposition, the filter paper and cotton ball were removed and each female was monitored daily until death. Wing length was recorded for all female mosquitoes to estimate fecundity for those mosquitoes whose fecundity was not directly measured (Armbruster and Hutchinson, 2002). All females’ wings were mounted on a glass side to measure the wing length from the distal end of the alula to the apex of the wing using a dissecting scope and micrometer.

### Life History Traits

We measured five traits relevant to population dynamics: larval survival, time to emergence, fecundity, adult longevity, and wing length (a standard proxy for fecundity (Armbruster and Hutchinson, 2002)). Larval survival was modeled as a binomial random variate for the number of larvae surviving from the 1^st^-instar larval stage until adult emergence. Time to emergence was measured as the median time for larvae to develop from 1^st^ instar to an adult per jar, in days. The median was used because the distribution of emergence times within a jar was right skewed. Fecundity was the number of eggs laid during the first oviposition event. Females that did not oviposit were assigned a fecundity of zero. Adult longevity was the number of days between adult emergence and death. Wing length was the distance from the distal end of the alula to the apex of the wing in mm. All traits were only measured for females, and the number of females per treatment at the start of the experiment was assumed to be 50% of the initial number of larvae within each jar.

We used generalized linear mixed models to test for the effect of temperature, *Ae. aegypti* density, *An. stephensi* density, and the interactions between temperature and species’ densities on each life history trait. We included replicate as a random intercept in all analyses. The models for larval survival were fit using a binomial distribution and a logit link. Because of the frequent occurrence of jars with no *An. stephensi* surviving, we used a hurdle model, which allows the degree of zero-inflation to vary across observations dependent on other predictor variables (Brooks et al., 2017). In our model, the structural zero-inflation term was dependent on both species’ densities. The time to emergence was modeled as the day that 50% of the females emerged and was fit with a generalized Poisson distribution and log link for both species. The generalized Poisson distribution is a mixture of Poisson distributions that is similar to a negative binomial distribution, but is more appropriate for right-skewed data due to its long tail (Joe and Zhu, 2005). Fecundity was modeled with a negative binomial distribution and log link, including a term for zero-inflation to account for some females that laid no eggs. Adult longevity was modeled with a generalized Poisson distribution and log link. Wing length was modeled with a gaussian distribution and identity link. Due to the low survival of *An. stephensi* larvae, sample size for this species was low, and models with interactions only converged when the response variable was the time to emergence or wing length. Therefore, the other three *An. stephensi* models included main effects only, as our data did not contain enough information to explain these interactions. All models were fit using the `glmmTMB` package (Brooks et al., 2017) and we assessed the residuals for divergence from normality using the `DHARMa` package (Hartig, 2019) in R v. 3.6.4 (R Core Team, 2018).

### Calculating the Per Capita Growth Rate

We calculated the per capita growth rate for each treatment following Chmielewski et al. (2010), substituting our own empirically-measured traits. We define the per capita growth rate, *r*, as the change in the population-level abundance of female mosquitoes attributable to one female mosquito:

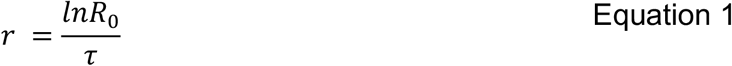

where *R*_*0*_ is the change in population size (ΔN) in one generation and τ is the generation time, or mean time to maturity and reproduction. *R*_*0*_ is the total population fecundity divided by the initial population size (*N*_*0*_):

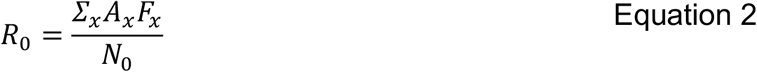

where *F*_*x*_ is the total lifetime reproduction of an individual emerging on day *x* and *A*_*x*_ is the number of individuals emerging on day *x*.

Lifetime fecundity, *F*_*x*_, is calculated from measured values of gonotrophic cycle length in days *(g)*, adult lifespan *(l)*, and the number of eggs per gonotrophic cycle (*f*_*x*_):

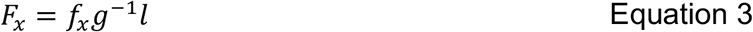

When *f*_*x*_ was not directly measured for an individual, it was approximated by a species-specific linear regression relating wing length (*w*_*x*_, in mm) to fecundity from experimental data (*Ae. aegypti*: *f*_*x*_ = −98.51 + 52.427*w*_*x*_, An, stephensi: *f*_*x*_ = −51.57 + 36.757*w*_*x*_). Some females had no intact wings to measure, and these were assigned the mean fecundity value for that temperature and density treatment for that species in that replicate.

Following Livdahl and Sugihara (1984), τ was weighted by the overall contribution to population fecundity:

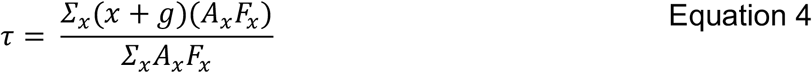

where *x* is the day of emergence, *g* is the gonotrophic cycle length in days, and *A*_*x*_ and *F*_*x*_ are the number of females that emerged on day *x* and their predicted lifetime fecundity, respectively.

This results in a final equation for the per capita growth rate for each density x temperature treatment:

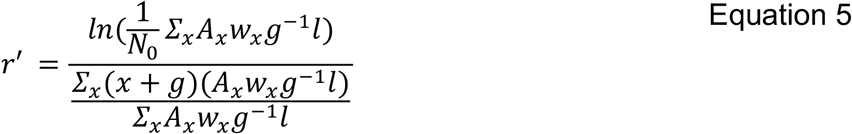

### Assessing the temperature-dependence of competition

As there is no consensus model for competitive interactions between mosquito species, we fit five theoretical discrete-time competition models that differ primarily in the shape of the response of the per capita growth rate to increasing species densities for each temperature treatment and mosquito species. We selected the best fit model using AIC (Table S2). All models were within 2 AIC of each other, and we chose the model with the lowest AIC, which approximates a Lotka-Volterra competition model where growth rates decline linearly with increasing species densities:

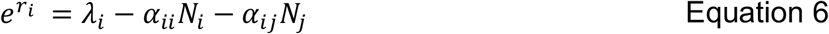

where *r*_*i*_ is the per capita growth rate of species *i*, *λ*_*i*_ is the intrinsic growth rate of species *i*, *α*_*ii*_ is the competition coefficient of intraspecific competition, *α*_*ij*_ is the competition coefficient of interspecific competition, and *N*_*i*_ and *N*_*j*_ are the starting population densities of species *i* and *j*. We parameterized this model separately for each temperature level and species to explore how λ, *α*_*ii*_, and *α*_*ij*_ changed as a function of the larval environment, and therefore if competitive interactions are temperature-dependent. We estimated the three parameters by fitting a non-linear least squares regression in R v. 3.6.3 (R Core Team, 2018).

We used the fit parameters from these equations to estimate each species’ competitive ability (*K*_i_) at each temperature following Hart et al. (2018):

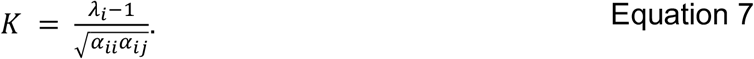

Briefly, this definition of competitive ability is derived from the concept of mutual invasibility, which states that each species must be able to invade while the other is at equilibrium in order for the species to coexist (Chesson, 2000). Solving for the conditions that allow for mutual invasion results in a ratio representing the average fitness difference of the two species, from which we derive each species fitness, or competitive ability, *K*. Importantly, this metric incorporates growth in the absence of competition (*λ*_*i*_ − 1) and the species ability to tolerate both intra- and inter-specific competition 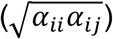. Because *K* incorporates a species’ growth rate and tolerance for competition, it allows for the possibility of a species with a low overall growth rate, but which is very robust to competition, to be the dominant long-term competitor. Thus, a species that is tolerant of competition, even it grows slowly, can persist and grow to high overall abundance, increasing the overall amount of competitive pressure and ultimately outcompeting less tolerant species regardless of their growth rates. By choosing to use these parameters, rather than direct measurements of fecundity, this definition of competitive ability incorporates differences across all life history stages to evaluate long-term competitive outcomes.

## Results

Our factorial design included fifteen species density combinations across five temperature treatments (75 treatments total) replicated three times. In one replicate, one jar (8 Ae. *aegypti*: 24 *An. stephensi* at 24°C) had only male Ae. *aegypti* emerge, and so this was dropped from all analyses, resulting in 179 jars across three replicates. After adult emergence, we followed a total of 1242 *Ae. aegypti* and 134 *An. stephensi* females to estimate fecundity and adult longevity.

### Life History Traits

Both species’ larval survival rates exhibited a unimodal response to temperature, with survival highest at intermediate temperatures (Fig. S1 A,D, Table 1, 2). This response was strongest for *Ae. aegypti* mosquitoes at high overall densities. In general, Ae. *aegypti* survival decreased with increasing intraspecific densities, although the shape of this relationship depended on temperature (Fig. S1B, Table 1). There was no evidence for an effect of interspecific densities on Ae. *aegypti* larval survival (Fig. S1C, Table 1). *An. stephensi* survival was strongly negatively impacted by both interspecific and intraspecific densities and no *An. stephensi* survived above interspecific densities of 32 (Fig. S1 E,F, Table 2).

**Table 1.** Table of coefficients from models of *Ae. aegypti* life history traits and per capita rates of change. 95% confidence intervals are below in parentheses. For larval survival thru longevity, if CIs overlap 1, it suggests that the predictor variable did not have a significant effect on the life history trait. Similarly, if estimates of the coefficients for wingspan and the per capita rate of change overlap 0, it suggests that the predictor variable did not have a significant effect on the life history trait. Significant effects are bolded.

| Predictors | Larval Survival | Time to Emergence | Fecundity | Longevity | Wing Length | Rate of Change |
| --- | --- | --- | --- | --- | --- | --- |
| Aedes Density | **0.67**<br>(0.62 - 0.72) | 1.01<br>(1.00 - 1.01) | **0.89**<br>(0.84 - 0.93) | 1.03<br>(1.00 - 1.07) | **-0.11**<br>(-0.12 - -0.10) | **-0.06**<br>(-0.11 - 0.00) |
| Anopheles Density | 0.99<br>(0.94 - 1.04) | 1.00<br>(1.00-1.00) | 1.01<br>(0.97 - 1.05) | 1.01<br>(0.99 - 1.04) | -0.01<br>(-0.02 - 0.00) | 0.00<br>(-0.05 - 0.04) |
| bs(Temperature)1 | 12.45<br>(0.95 - 163.39) | **0.36**<br>(0.31 - 0.43) | 1.17<br>(0.37 - 3.68) | **3.16**<br>(1.45 - 6.88) | 0.06<br>(-0.24 - 0.37) | 0.87<br>(-0.19 - 1.93) |
| bs(Temperature)2 | **28.94**<br>(2.49 - 336.02) | **0.28**<br>(0.23- 0.33) | 1.98<br>(0.84 - 4.71) | 0.58<br>(0.31 - 1.07) | -0.26<br>(-0.52 - 0.00) | **1.11**<br>(0.46 - 1.76) |
| bs(Temperature)3 | **4.17**<br>(1.50 - 11.59) | **0.30**<br>(0.28-0.33) | 0.62<br>(0.37 - 1.04) | 1.21<br>(0.84 - 1.74) | **-0.67**<br>(-0.81 - -0.53) | **0.76**<br>(0.28 - 1.25) |
| Aedes Density x<br>bs(Temperature)1 | 0.83<br>(0.64 - 1.09) | 1.02<br>(1.00-1.04) | 0.96<br>(0.84 - 1.11) | 0.95<br>(0.86 - 1.04) | -0.02<br>(-0.05 - 0.02) | -0.03<br>(-0.16 - 0.11) |
| Aedes Density x<br>bs(Temperature)2 | 0.83<br>(.64 - 1.06) | 1.01<br>(0.99 - 1.03) | 0.96<br>(0.86 - 1.07) | 1.08<br>(1.00 - 1.16) | 0.00<br>(-0.03 - 0.03) | -0.01<br>(-0.10 - 0.07) |
| Aedes Density x<br>bs(Temperature)3 | **0.84**<br>(0.75 - 0.94) | 1.00<br>(0.99-1.01) | 1.01<br>(0.95 - 1.08) | 0.97<br>(0.94 - 1.01) | **0.02**<br>(0.01 - 0.04) | -0.01<br>(-0.07 - 0.05) |
| Anopheles Density x<br>bs(Temperature)1 | 1.02<br>(0.86 - 1.20) | 1.01<br>(0.99 -1.02) | 1.01<br>(0.92 - 1.11) | 0.96<br>(0.90 - 1.03) | -0.01<br>(-0.03 - 0.02) | -0.01<br>(-0.11 - 0.10) |
| Anopheles Density x<br>bs(Temperature)2 | 1.04<br>(0.89 - 1.21) | 1.00<br>(0.99 -1.02) | **0.93**<br>(0.86 - 0.99) | 1.04<br>(0.98 - 1.09) | 0.01<br>(-0.01 - 0.03) | 0.01<br>(-0.05 - 0.08) |
| Anopheles Density x<br>bs(Temperature)3 | **1.08**<br>(1.01 - 1.16) | 1.00<br>(1.00-1.01) | 0.99<br>(0.95 - 1.04) | 0.99<br>(0.96 - 1.02) | 0.00<br>(-0.01 - 0.01) | 0.00<br>(-0.05 - 0.05) |
| Zero-Inflation Intercept |  |  | 0.28<br>(0.24 - 0.32) |  |  |  |
| Observations | 179 | 179 | 1242 | 1194 | 1088 | 174 |
| Marginal / Conditional R2 | 0.371 / 0.387 | 0.987 / 0.989 | 0.175 / 0.204 | 0.093 / 0.208 | 0.736 / 0.741 | 0.925 / 0.969 |

**Table 2.** Table of coefficients from models of *An. stephensi* life history traits and per capita rates of change. 95% confidence intervals are below in parentheses. For larval survival thru longevity, if CIs overlap 1, it suggests that the predictor variable did not have a significant effect on the life history trait. Similarly, if estimates of the coefficients for wingspan and the per capita rate of change overlap 0, it suggests that the predictor variable did not have a significant effect on the life history trait. Significant effects are bolded.

| Predictors | Larval Survival | Time to Emergence | Fecundity | Longevity | Wing Length | Rate of Change |
| --- | --- | --- | --- | --- | --- | --- |
| **Aedes Density** | **0.82**<br>(0.71 - 0.95) | 0.99<br>(0.95 - 1.04) | 0.94<br>(0.80 - 1.12) | 0.99<br>(0.92 - 1.06) | 0.03<br>(-0.07 - 0.13) | 0.00<br>(0.00 - 0.01) |
| **Anopheles Density** | **0.62**<br>(0.53 - 0.72) | 0.99<br>(0.95 - 1.03) | 0.94<br>(0.70 - 1.25) | 0.95<br>(0.88 - 1.02) | 0.09<br>(-0.07 - 0.25) | 0.00<br>(0.00 - 0.00) |
| **bs(Temperature)1** | 13.42<br>(2.50 - 72.09) | **0.17**<br>(0.04 - 0.71) | 1.15<br>(0.52 - 2.51) | 1.92<br>(0.83 - 4.47) | **2.71**<br>(0.46 - 4.96) | **0.20**<br>(0.11 - 0.28) |
| **bs(Temperature)2** | 0.79<br>(0.24 - 2.61) | 0.55<br>(0.03 - 10.87) | - | 1.08<br>(0.59 - 1.97) | -2.10<br>(-4.65 - 0.44) | - |
| **bs(Temperature)3** | 1.57<br>(0.64 - 2.83) | 0.55<br>(0.07 - 4.33) | - | 1.23<br>(0.80 - 1.89) | 0.79<br>(-1.17 - 2.76) | - |
| **Aedes Density x**<br>**bs(Temperature)1** | - | 1.19<br>(0.99 - 1.42) | - | - | -0.21<br>(-0.46 - 0.04) | - |
| **Aedes Density x**<br>**bs(Temperature)2** | - | 0.91<br>(0.67 - 1.23) | - | - | 0.06<br>(-0.18 - 0.31) | - |
| **Aedes Density x**<br>**bs(Temperature)3** | - | 0.95<br>(0.79 - 1.15) | - | - | -0.10<br>(-0.27 - 0.07) | - |
| **Anopheles Density x**<br>**bs(Temperature)1** | - | 1.14<br>(0.93 - 1.40) | - | - | **-0.46**<br>(-0.82 - -0.09) | - |
| **Anopheles Density x**<br>**bs(Temperature)2** | - | 0.94<br>(0.58 - 1.53) | - | - | 0.21<br>(-0.20 - 0.63) | - |
| **Anopheles Density x**<br>**bs(Temperature)3** | - | 0.90<br>(0.64 - 1.28) | - | - | -0.25<br>(-0.59 - 0.09) | - |
| **Zero-Inflation Intercept** | 0.00<br>(0.00 - 0.02) | - | 2.83<br>(1.92 - 4.16) | - | - | - |
| **Zero-Inflation**<br>**Aedes Density** | 2.35<br>(1.70 - 3.26) | - | - | - | - | - |
| **Zero-Inflation**<br>**Anopheles Density** | 2.14<br>(1.57 - 2.92) | - | - | - | - | - |
| **Observations** | 180 | 52 | 134 | 127 | 127 | 23 |
| **Marginal/ Conditional R2** | 0.304 / 0.328 | 0.896 / 0.909 | 0.547 / 0.641 | 0.046 / 0.09 | 0.547 / 0.641 | 0.508 / 0.657 |

Both species’ emerged more quickly with increasing temperatures and neither species’ time to emerge was impacted by interspecific or intraspecific densities (Fig. S2, Table 1, 2). However, the time to emergence was generally longer for *An. stephensi* larvae than Ae. *aegypti* larvae at all treatment combinations (Fig. S2).

There was no evidence for an effect of temperature on Ae. *aegypti* female fecundity (Fig. S3A, Table 1). Higher intraspecific densities resulted in lower fecundity (Fig. S3B, Table 1) regardless of temperature treatment, but interspecific density had no effect (Fig. S3C, Table 1). We found no evidence for an effect of temperature, interspecific density, or intraspecific density on *An. stephensi* fecundity (Fig. S3 E,F, Table 2).

Ae. *aegypti* longevity was highest in females that were reared at intermediate temperatures (Fig. S4A, Table 1). We failed to find evidence for an effect of intraspecific or interspecific densities on *Ae. aegypti* longevity (Fig. S4B,C, Table 1). We found no evidence for a difference in *An. stephensi* longevity across any of the three treatments (Fig. S4 D,E,F, Table 2). Across all temperatures, *Ae. aegypti* females lived approximately twice as long as *An. stephensi* females, with mean adult lifespans of 32.1 ± 16.0 *sd* and 16.7 ± 9.80 *sd* days, respectively.

Increasing temperatures led to shorter wing lengths for both species (Fig. S5 A,D, Table 1, 2). Ae. *aegypti* wing lengths also decreased with increasing intraspecific densities (Fig. S5 B, Table 1), but were unaffected by interspecific densities (Fig. S5 C, Table S). Neither species’ density was found to influence *An. stephensi* wing lengths (Fig. S5 E,F, Table 2).

### Per Capita Rate of Change

Temperature significantly influenced both species’ per capita rate of change. Ae. *aegypti* growth rates had a unimodal relationship with temperature (Fig. S6A, Table 1) and *An. stephensi* growth rates increased with increasing temperatures (Fig. S6D, Table 2). Ae. *aegypti* growth rates decreased with increasing Ae. *aegypti* densities (Fig. S6B, Table 1), but we found no evidence for an effect of *An. stephensi* density on Ae. *aegypti* growth rates (Fig. S6C, Table 1). We found no support for an effect of either species’ densities on *An. stephensi* growth rates (Fig. S6 E,F, Table 2).

### Temperature-Dependence of Competition

The parameters from fitted competition models differed across temperature levels (Fig. 2). Both species’ intrinsic growth rates (λ) were dependent on temperature (Fig. 2A). Ae. *aegypti* growth rates had a unimodal relationship with temperature, peaking at 28°C. *An. stephensi* growth rates increased with temperature (Fig. 2). However, competition models did not converge for *An. stephensi* at 24°C and 32°C due to a low sample size, rendering the shape of *An. stephensi* ‘s response to temperature less clear. Both species’ intraspecific competition coefficients (αii) increased with temperature (Fig. 2B). The effect of *Ae. aegypti* density on *An. stephensi* population growth rates increased with increasing temperatures, while the effect of *An. stephensi* did not differ from zero at any temperature. Inserting these parameters into the general equation for competitive ability (Eq. 7) illustrates that *Ae. aegypti* is the dominant competitor at all temperatures tested (Fig. 2D).

**Figure 2.**
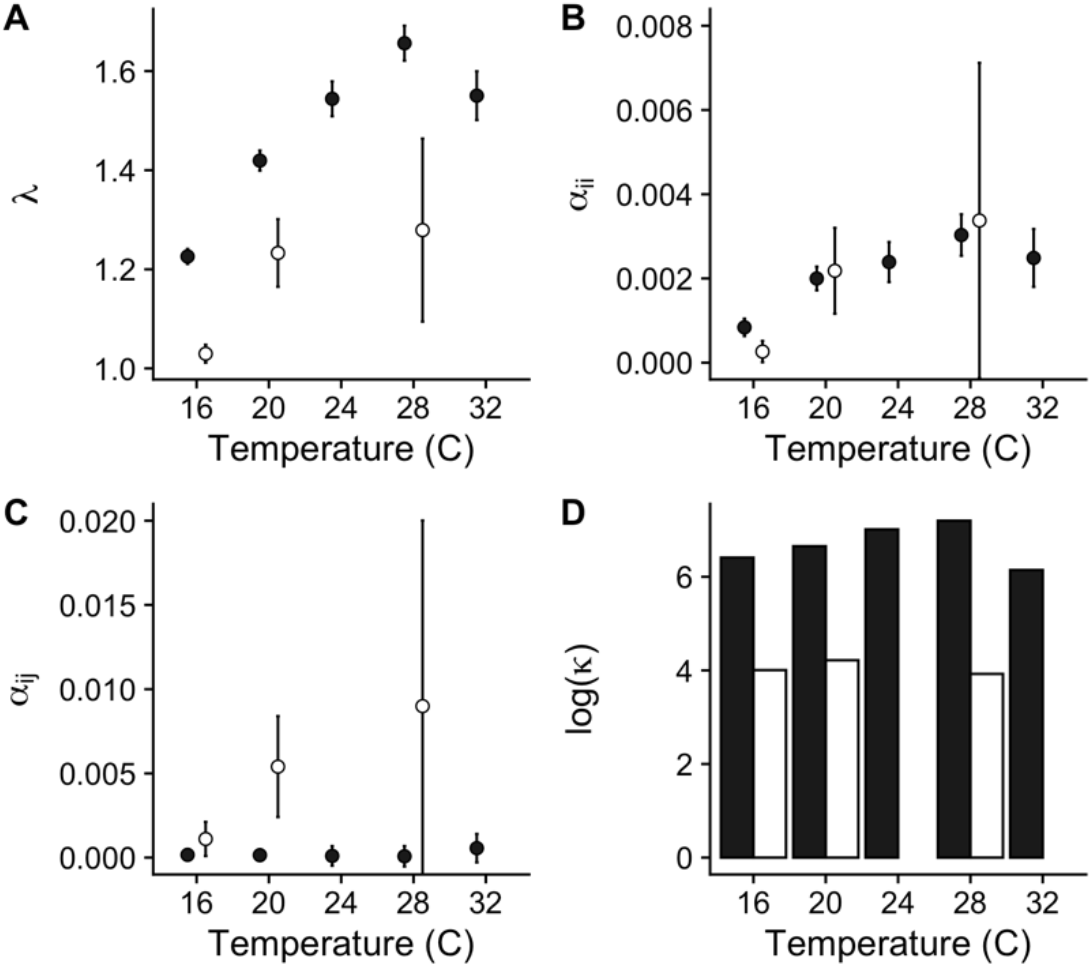
Parameter estimates from the competition model for Ae. *aegypti* (filled) and *An. stephensi* (unfilled). Panels represent. A) intrinsic growth rates (λ_i_), B) the intraspecific competition coefficient α_ii_, C) the interspecific competition coefficient α_ij_, and D) a comparison of species competitive abilities (on a natural-log scale). Sample sizes were too low at 24°C and 32°C to estimate parameters for *An. stephensi*.

## Discussion

Mosquito vectors are rapidly expanding their ranges into new geographic areas and competing with resident mosquito species (e.g. *Ae. albopictus* in the US and Europe, *Ae. koreicus* in Italy (Marcantonio et al., 2016), *Cx. coronator* in Southeastern US (Wilke et al., 2020)). Understanding how novel interactions may affect mosquito population dynamics is necessary for predicting disease risk at invasion fronts, such as that of *An. stephensi* (Seyfarth et al., 2019; Takken and Lindsay, 2019). We found strong evidence for asymmetric competition between *Ae. aegypti* and *An. stephensi* across the full range of temperatures tested in this experiment, with *Ae. aegypti* consistently emerging as the dominant competitor. The intrinsic growth rate of *An. stephensi* was lower than that of *Ae. aegypti* at all temperatures, and *An. stephensi* was less tolerant of interspecific competition than *Ae. aegypti*. Given the global range of *Ae. aegypti,* this competitive interaction has the potential to influence the rate of spread of *An. stephensi* in its invasive range.

Life-history traits were influenced by both temperature and species densities. Ecothermic metabolic theory predicts that colder temperatures result in longer development times that translate into larger bodied female mosquitoes that have higher fecundity rates (Kingsolver and Huey, 2008). This has often been the case in mosquito systems (Armbruster and Hutchinson, 2002 but see Reiskind and Zarrabi, 2012). In our study, colder temperatures resulted in longer development times and wing lengths for both species, but the effect of temperature on fecundity was much weaker. The size-fecundity relationship in mosquitoes is typically weakest towards the thermal minima (Costanzo et al., 2018), with fecundity saturating with decreasing temperatures. In this study, measured fecundity at 16°C and 20°C was less than predicted by a linear size-fecundity relationship (Fig. S3A), mirroring this breakdown of the relationship at cold extremes. Interestingly, we did not find evidence that species’ densities influenced either species’ time to emergence, in contrast with previous studies on larval competition and mosquito development times (Couret et al., 2014). However, Ae. *aegypti* wing length and fecundity did decrease with increasing conspecific densities. This is in agreement with other studies that found that limiting resources via competition leads to smaller bodied mosquitoes (Alto et al., 2005; Juliano et al., 2014). Rather than delaying emergence to develop into larger bodied mosquitoes given resource limitations, mosquitoes under higher competitive pressures in our study emerged at similar times, but with smaller bodies.

While we found that both species’ intrinsic growth rates and competition coefficients were temperature-dependent, this difference did not affect the outcome of competition. The species’ responses to conspecific competition were comparable, but the effect of *Ae. aegypti* on *An. stephensi* was much stronger than the effect of *An. stephensi* on *Ae. aegypti*. In the field, both species breed in artificial containers, but *Ae. aegypti* is more tolerant to overcrowding than *An. stephensi* (Yadav et al., 2017). This agrees with our finding that larval *An. stephensi* survival was low at high densities. *Ae. aegypti*’s tolerance of crowding may be a result of the close association of *Ae. aegypti* with humans throughout its evolutionary history, leading to adaptations well-suited for human-modified landscapes (Brown et al., 2014). Adult *An. stephensi* were generally larger and had a longer development period than *Ae. aegypti* in our study. This concurs with allometric theory, which predicts a positive relationship between organism body size and development time for ectotherms (Gillooly et al., 2002). This difference in larval development rates led to a longer generational period and lower intrinsic growth rate for *An. stephensi* compared to *Ae. aegypti,* contributing to differences in competitive ability.

Our laboratory experiment suggests that long term coexistence between the two species is unlikely; yet, the two species co-exist at the landscape-scale in the endemic range of *An. stephensi* and *An. stephensi* is currently expanding into the range of *Ae. aegypti.* Our study only considers interactions within one larval habitat and does not account for mechanisms that act at a larger-scale that could explain the co-existence patterns found in the field. One such mechanism is the classical competition-dispersal trade-off: metacommunity level coexistence is possible if the inferior competitor’s superior dispersal ability allows it to colonize new patches where the competitive pressure is lower (Hastings, 1980). Indeed, *Ae. aegypti* flight range is estimated to be 83.4 ± 52.2m, compared to a longer dispersal distance of 144.5 ± 53.0m for *An. stephensi* (Verdonschot and Besse-Lototskaya, 2014). Thus, while the species may not coexist within a single larval habitat, the wider dispersal range of *An. stephensi* may allow for landscape-scale coexistence at the level of the metacommunity. Similarly, species-specific microhabitat preferences may reduce the frequency of habitat overlap, and thereby competition, during the larval stage. While *An. stephensi* and *Ae. aegypti* are found together in small artificial containers, *An. stephensi* also oviposit in larger water bodies, such as overhead water tanks, where *Ae. aegypti* is less common (Thomas et al., 2016). These water bodies may serve as a refugia for *An. stephensi* from high competitive pressure by *Ae. aegypti* and allow for broad-scale coexistence. Species differences at non-competing life stages may also allow for coexistence in the field. For example, our models assumed an equal embryonic development period for both species between oviposition and hatching. No studies have directly measured *An. stephensi* embryonic development times, but a study in *An. albitarsis* estimate that it takes around 34hrs for eggs to fully mature, noting development timings similar to *An. maculipennis* (Monnerat et al., 2002). In comparison, *Ae. aegypti* eggs are fully developed around 90hrs and require a week for embryonation (Raminani and Cupp, 1978). These species differences in egg maturation rates could lengthen our estimated generation times for *Ae. aegypti,* decreasing the species’ intrinsic growth rates. Finally, interpopulation differences in competitive ability have been found in *Ae. aegypti* (Leisnham and Juliano, 2010), and invasive *An. stephensi* populations may have an increased competitive ability compared to those in endemic areas (e.g. the Evolution of Increased Competitive Ability hypothesis (Strayer et al., 2006)). If the invader genotype of *An. stephensi* is able to coexist with, or even outcompete, *Ae. aegypti*, then the result of strong asymmetric competition we found with our specific populations may not apply to competition in the region of North Africa where *An. stephensi* is currently invading. Further research that includes a variety of field-derived genotypes in competition experiments could explore the importance of genotype x genotype interactions in this system.

While we found the difference in competitive ability between the two species to be strong across all temperatures, this could also be an artefact of our specific abiotic and biotic experimental conditions. Food type and availability alters the outcome of competition between other container mosquitoes (Juliano, 2010; Murrell and Juliano, 2008; Yee et al., 2007), and could play a role in this system as well. Other abiotic factors, such as desiccation tolerance of eggs (Juliano et al., 2002) or climate factors during adult life stages, may also change the outcome of this competitive interaction. The presence of additional species, such as predators (Juliano, 2009), other competitors (Bowden, 2016), or parasites (Westby et al., 2019), also have the potential to alter this pair-wise interaction. Additionally, the two strains used in this experiment have different domestication histories due to constraints on strain availability. The *Ae. aegypti* strain was recently derived from Mexico, while the *An. stephensi* (Liston) strain was originally established from an Indian population several decades ago and since kept in laboratory conditions. A laboratory strain could exhibit a high tolerance for crowding (Kesavaraju et al., 2012), but may also have reduced overall fitness due to inbreeding (Koenraadt et al., 2010). Additionally, these strains are not sympatric in nature, having originated in Mexico and India, and this combination may not reflect the competitive interactions between *Ae. aegypti* and *An. stephensi* in North Africa. Therefore, the overall effects of strain differences on competition outcomes are difficult to predict, but may alter our predictions regarding long-term coexistence.

Generally, invasive mosquito species exhibit a higher competitive ability than native species, leading to a reduction in the native population abundance or range (Juliano and Lounibos, 2005; Kaufman and Fonseca, 2014). Although *An. stephensi* is spreading into North Africa (Surendran et al., 2019), it may not significantly reduce the population abundance of existing *Ae. aegypti* populations as has been seen in the Southeastern US, given its low competitive ability relative to *Ae. aegypti.* Further, If *Ae. aegypti* is limiting the spread of *An. stephensi* into urban areas in the Middle East and North Africa, vector control efforts targeting *Ae. aegypti* could have unexpected consequences for *An. stephensi* dynamics. For example, vector control efforts that reduce *Ae. aegypti* abundances could lead to an increase in *An. stephensi* abundances via competitive release. This unexpected consequence was observed when *Ae. albopictus* invaded urban centers in Manila, Philippines following a reduction of its competitor *Ae. aegypti* due to targeted insecticide spraying (Gilotra et al., 1967; Lounibos, 2007). Larval source reduction campaigns, however, could limit the abundance of both species by reducing their shared habitat. In areas with both mosquito vectors, vector control efforts should include a variety of approaches, rather than ones that target a single species, to avoid the competitive release of other mosquito vectors.

We found evidence for strong asymmetric competition between *Ae. aegypti* and *An. stephensi*. This study is the first test of competition between these two species and only considered one strain of each species across one environmental gradient (e.g. temperature). While more work is needed to assess the applicability of our results to the field, our study does suggest that competition is an important factor to consider in the context of the expanding range of *An. stephensi.* Temperature-based models of *An. stephensi* distribution predict high environmental suitability for the species in the Horn of Africa (Miazgowicz et al., 2019), but biotic interactions, such as competition, may alter this prediction. In addition to *An. stephensi,* many other mosquito species of public health importance have been the focus of temperature-based suitability models that rarely include biotic interactions (Golding et al., 2015). For *Ae. aegypti* and *Ae. albopictus*, especially, biotic interactions are hypothesized to limit species ranges (Lounibos and Juliano, 2018). Our study suggests that species interactions are also important for population dynamics of *An. stephensi*, and that these interactions should be empirically tested and, when appropriate, incorporated into our predictions of mosquito species ranges and invasion dynamics.

## DATA AVAILABILITY

All code and data needed to reproduce this analysis can be found at FIGSHARE URL.

## ACKNOWLEDGEMENTS

We would like to thank Justine Shiau for her help in the insectary and Paige Miller for her editorial feedback. This work was supported by the National Science Foundation Graduate Research Fellowship, National Science Foundation Research Experiences for Undergraduates (Grant No. 1156707), and the National Institutes of Health (Grant No. 5R01AI110793-04).

## SUPPLEMENTAL MATERIALS

### SUPPLEMENTAL FIGURES

**Figure S1.**
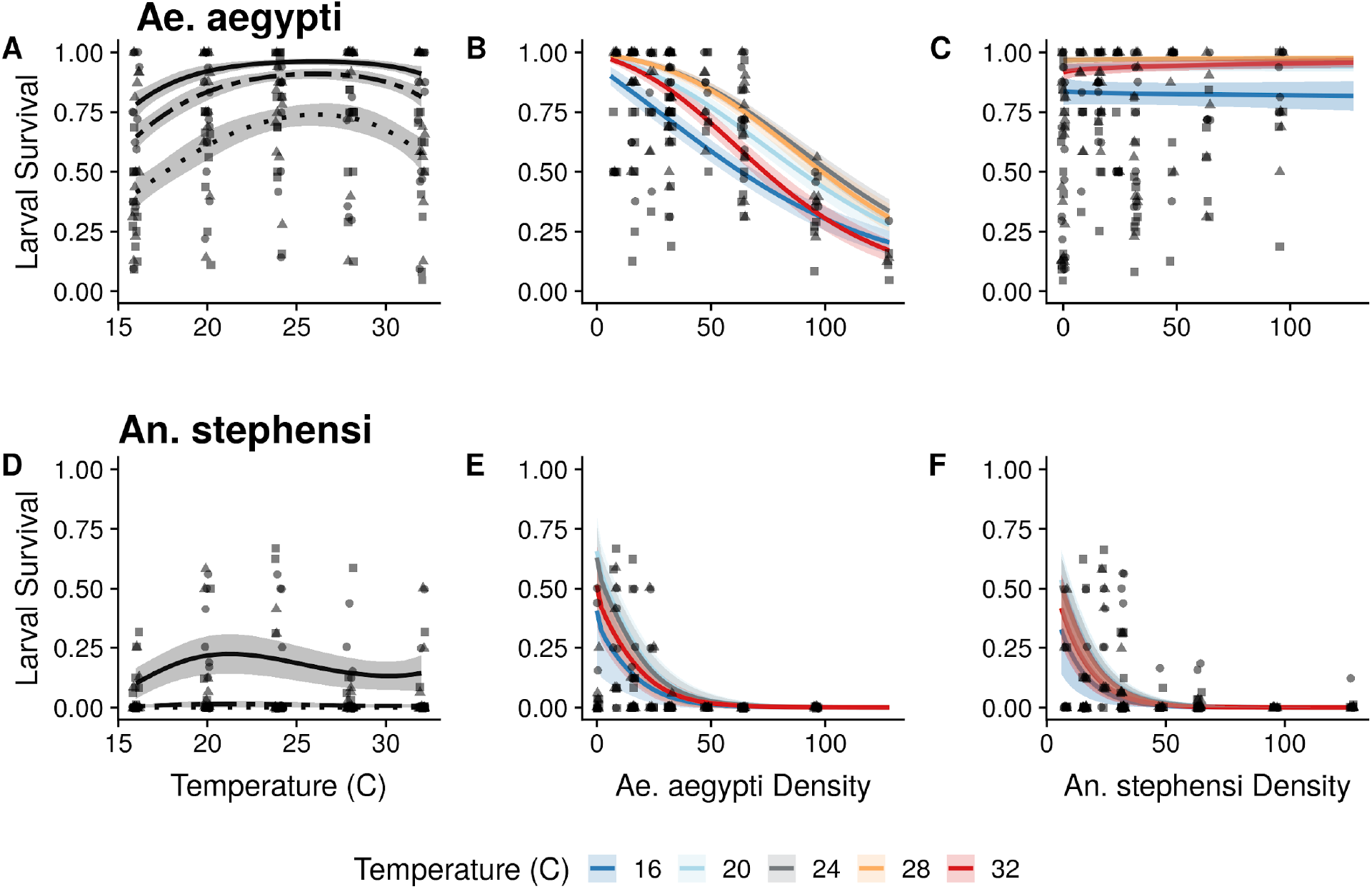
Effect of temperature (A,D), Ae. *aegypti* density (B,E) and *An. stephensi* density (C,F) on each species’ larval survival. Top row (A-C) is Ae. *aegypti* and bottom row (D-F) is *An. stephensi*. The points represent raw data, with each replicate denoted by a different symbol, and solid lines represent model fits with 95% CI. In panels A and D, three lies are shown for three unique species ratios, 16:16 (solid), 32:32 (dashed), 64:64 (dotted). In panels B,C and E,F, solid lines represent the model fit with the other species density held constant at 16.

**Figure S2.**
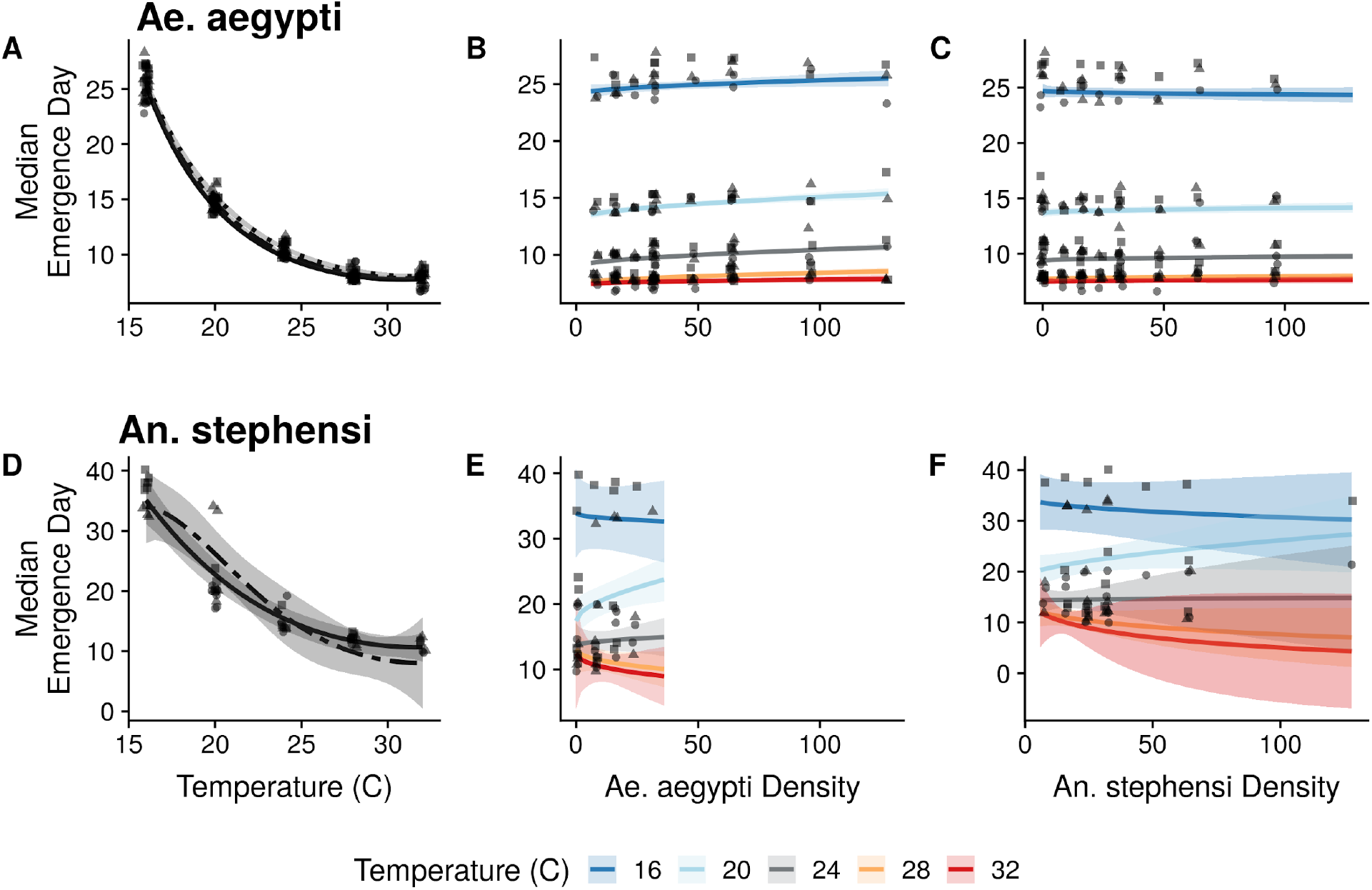
Effect of temperature (A,D),Ae. *aegypti* density (B,E) and *An. stephensi* density (C,F) on each species’ time to emergence. Top row (A-C) is Ae. *aegypti* and bottom row (D-F) is *An. stephensi*. The points represent raw data, with each replicate denoted by a different symbol, and solid lines represent model fits with 95% CI. In panels A and D, three lies are shown for three unique species ratios, 16:16 (solid), 32:32 (dashed), 64:64 (dotted). In panels B,C and E,F, solid lines represent the model fit with the other species density held constant at 16.

**Figure S3.**
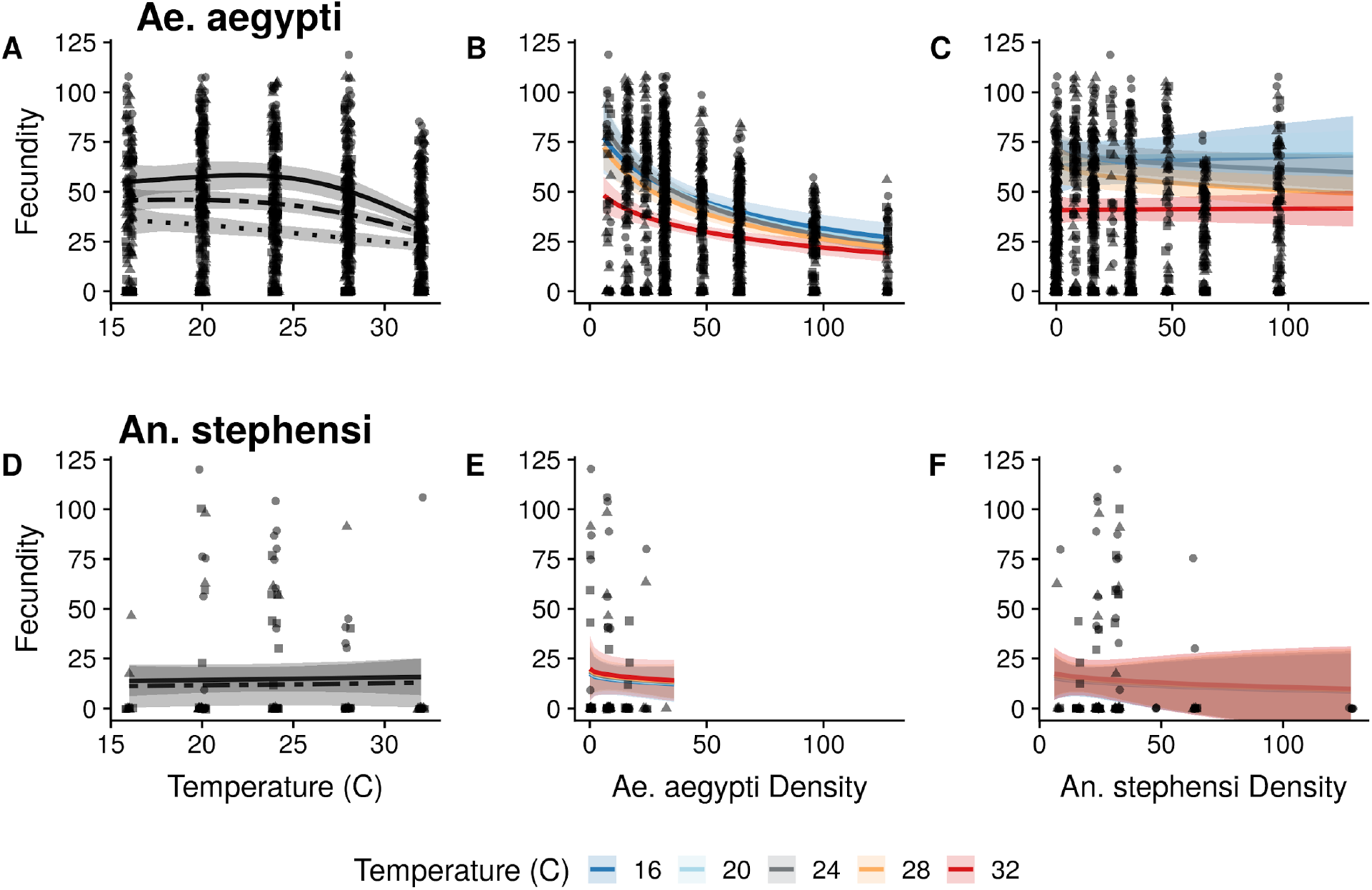
Effect of temperature (A,D), Ae. *aegypti* density (B,E) and *An. stephensi* density (C,F) on each species’ fecundity. Top row (A-C) is Ae. *aegypti* and bottom row (D-F) is *Anopheles*. The points represent raw data, with each replicate denoted by a different symbol, and solid lines represent model fits with 95% CI. In panels A and D, three lies are shown for three unique species ratios, 16:16 (solid), 32:32 (dashed), 64:64 (dotted). In panels B,C and E,F, solid lines represent the model fit with the other species density held constant at 16.

**Figure S4.**
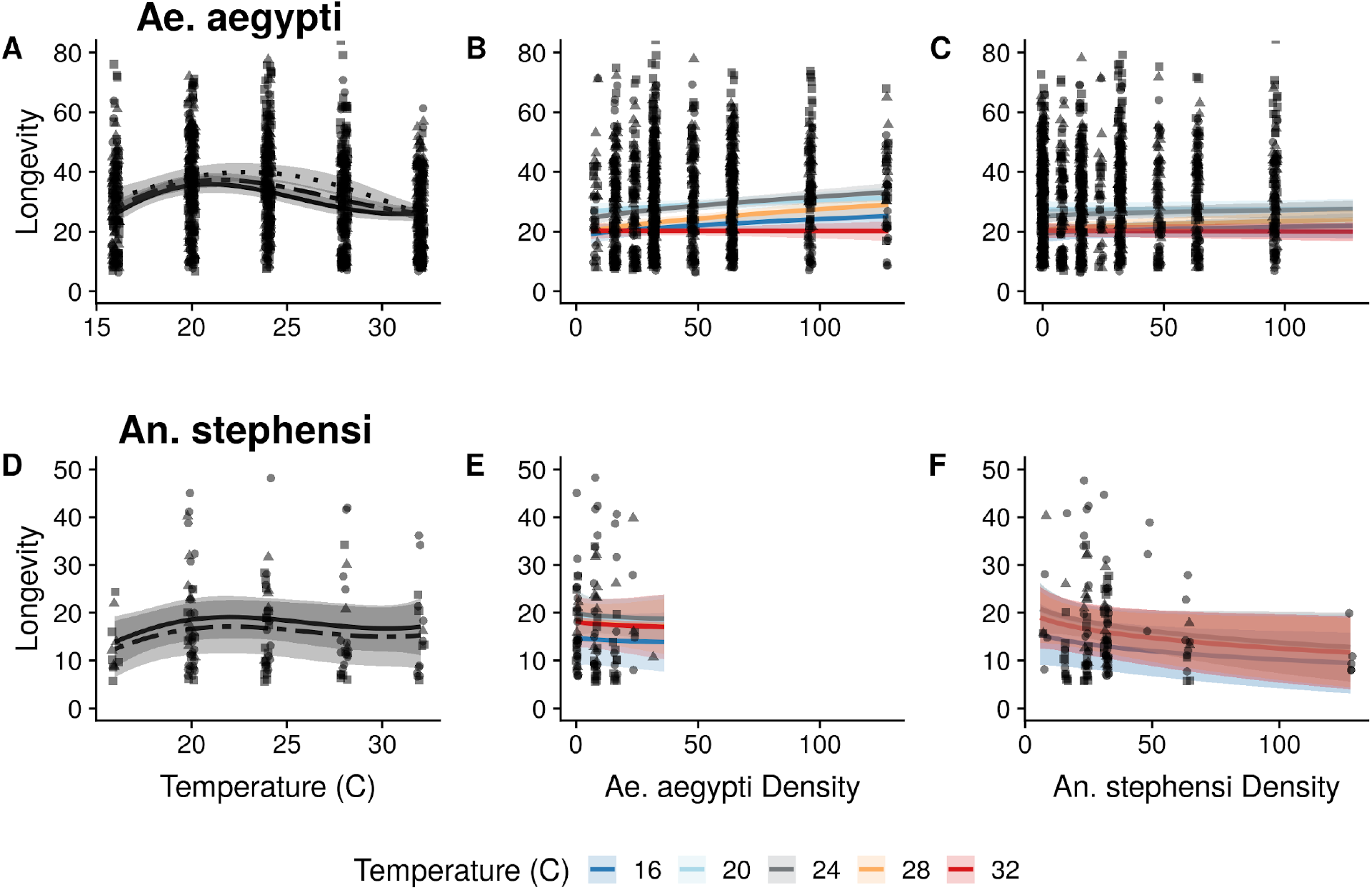
Effect of temperature (A,D), Ae. *aegypti* density (B,E) and *An. stephensi* density (C,F) on each species’ longevity. Top row (A-C) is Ae. *aegypti* and bottom row (D-F) is *Anopheles*. The points represent raw data, with each replicate denoted by a different symbol, and solid lines represent model fits with 95% CI. In panels A and D, three lies are shown for three unique species ratios, 16:16 (solid), 32:32 (dashed), 64:64 (dotted). In panels B,C and E,F, solid lines represent the model fit with the other species density held constant at 16.

**Figure S5.**
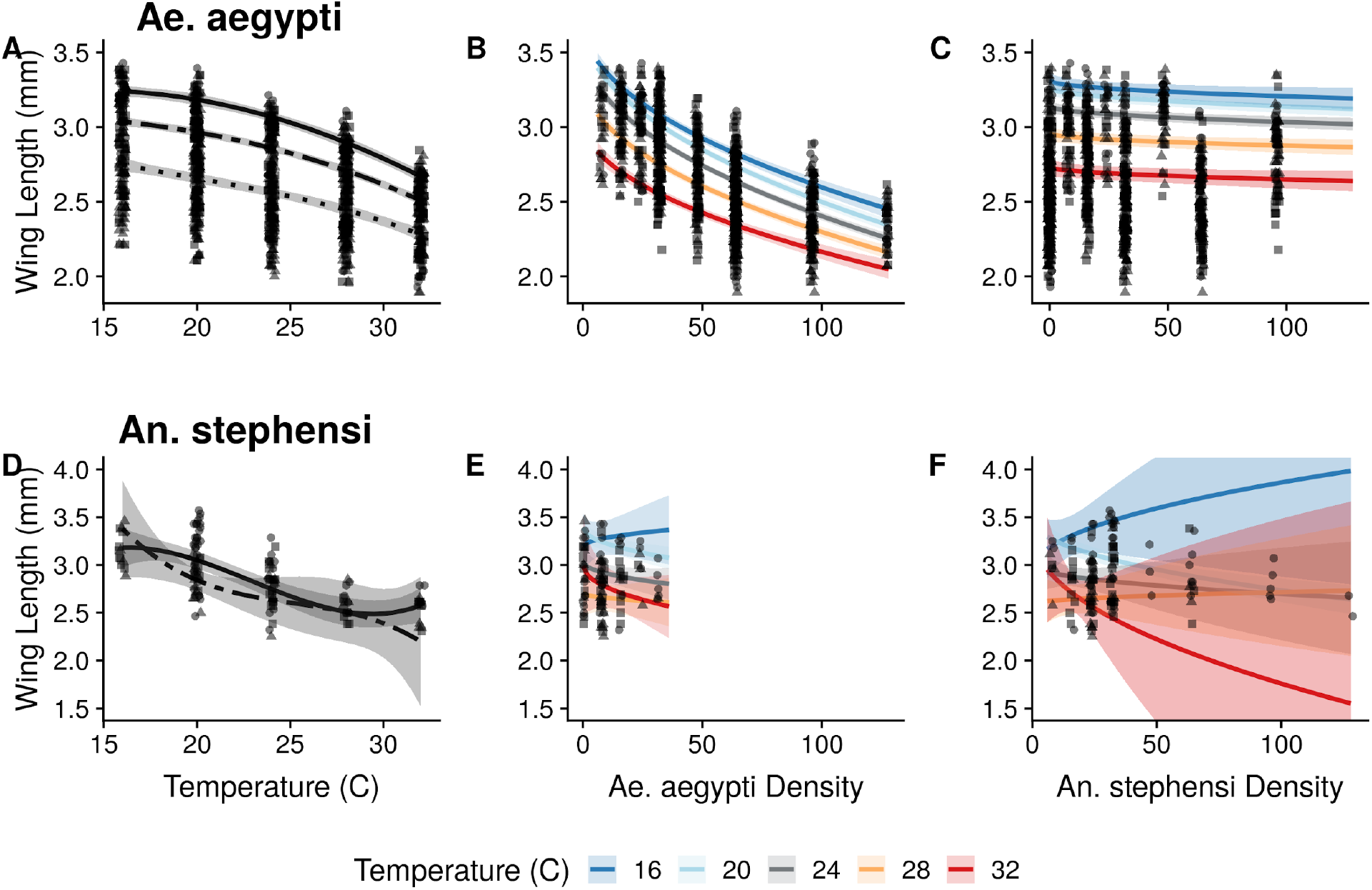
Effect of temperature (A,D), Ae. *aegypti* density (B,E) and *An. stephensi* density (C,F) on each species’ wing length. Top row (A-C) is Ae. *aegypti* and bottom row (D-F) is *An. stephensi*. The points represent raw data, with each replicate denoted by a different symbol, and solid lines represent model fits with 95% CI. In panels A and D, three lies are shown for three unique species ratios, 16:16 (solid), 32:32 (dashed), 64:64 (dotted). In panels B,C and E,F, solid lines represent the model fit with the other species density held constant at 16.

**Figure S6.**
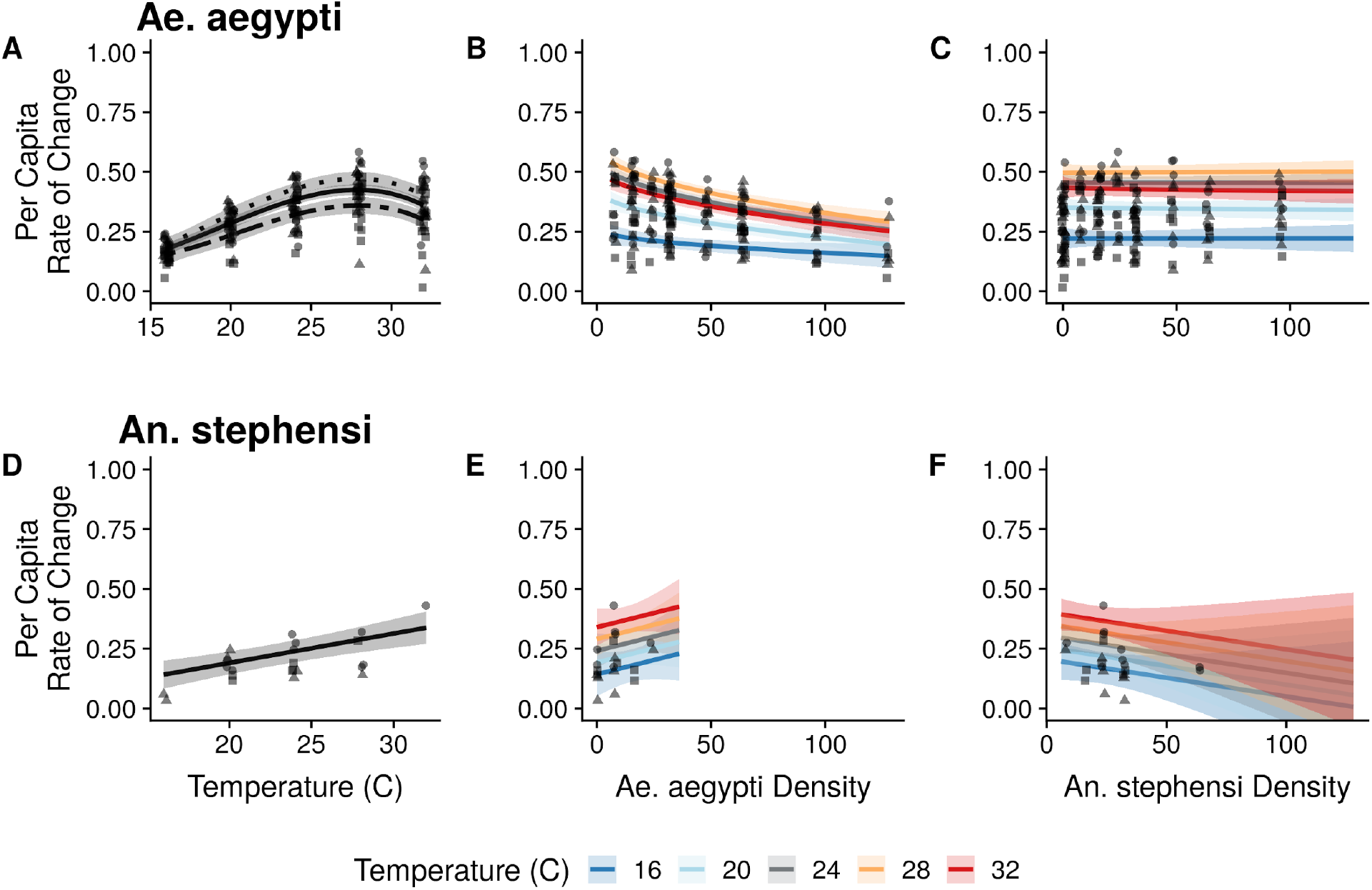
Effect of temperature (A,D), Ae. *aegypti* density (B,E) and *An. stephensi* density (C,F) on each species’ per capita rate of change. Top row (A-C) is Ae. *aegypti* and bottom row (D-F) is *An. stephensi*. The points represent raw data, with each replicate denoted by a different symbol, and solid lines represent model fits with 95% CI. In panels A and D, three lies are shown for three unique species ratios, 16:16 (solid), 32:32 (dashed), 64:64 (dotted). In panels B,C and E,F, solid lines represent the model fit with the other species density held constant at 16.

### SUPPLEMENTAL TABLES

**Table S1.** Species densities (per 250 mL) used in response surface design.

| *An. stephensi* | *Ae. aegypti* |
| --- | --- |
| 32 | 0 |
| 24 | 8 |
| 16 | 16 |
| 8 | 24 |
| 0 | 32 |
| 64 | 0 |
| 48 | 16 |
| 32 | 32 |
| 16 | 48 |
| 0 | 64 |
| 128 | 0 |
| 96 | 32 |
| 64 | 64 |
| 32 | 96 |
| 0 | 128 |

**Table S2.** AIC values resulting from fitting five theoretical competition models to our estimates of per capita growth rates for *Ae. aegypti λ*_*i*_, *α*_*ii*_, and *α*_*ij*_ are all temperature-dependent.

| Model | df | AIC |
| --- | --- | --- |
| $e^{r_i} = \frac{\lambda_i}{1 + \alpha_{ii} N_i + \alpha_{ij} N_j}$ | 16 | -314.3939 |
| $e^{r_i} = \frac{\lambda_i}{(1 + \alpha_{ii} N_i + \alpha_{ij} N_j)^b}$ | 17 | -313.2340 |
| $e^{r_i} = \lambda_i e^{-\alpha_{ii} N_i - \alpha_{ij} N_j}$ | 16 | -314.8702 |
| $e^{r_i} = \frac{\lambda_i}{1 + N_i^{\alpha_{ii}} + N_j^{\alpha_{ij}}}$ | Unable to fit | Unable to fit |
| $e^{r_i} = \lambda_i - \alpha_{ii} N_i - \alpha_{ij} N_j$ | 16 | -315.1535 |

## References

Alto, B.W., Lounibos, L.P., Higgs, S., Juliano, S.A., 2005. Larval competition differentially affects arbovirus infection in Aedes mosquitoes. Ecology 86, 3279–3288.

Armbruster, P., Hutchinson, R.A., 2002. Pupal mass and wing length as indicators of fecundity in Aedes albopictus and Aedes geniculatus (Diptera: Culicidae). J Med Entomol 39, 699–704. https://doi.org/10.1603/0022-2585-39.4.699

Bowden, S.E., 2016. Trans-boundary ecosystem effects on vector community diversity: implications for dilution and amplification in multi-species host-pathogen systems (Doctoral Dissertation). University of Georgia, Athens, GA.

Brady, O.J., Hay, S.I., 2020. The Global Expansion of Dengue: How Aedes aegypti Mosquitoes Enabled the First Pandemic Arbovirus. Annual Review of Entomology 65, null. https://doi.org/10.1146/annurev-ento-011019-024918

Brooks, M.E., Kristensen, K., van Bentehm, K.J., Magnusson, A., Berg, C.W., Nielsen, A., Skaug, H.J., Maechler, M., Bolker, B.M., 2017. glmmTMB Balances Speed and Flexibility Among Packages for Zero-inflated Generalized Linear Mixed Modeling. The R Journal 9, 378–400.

Brown, J.E., Evans, B.R., Zheng, W., Obas, V., Barrera-Martinez, L., Egizi, A., Zhao, H., Caccone, A., Powell, J.R., 2014. Human impacts have shaped historical and recent evolution in Aedes aegypti, the dengue and yellow fever mosquito. Evolution 68, 514–525. https://doi.org/10.1111/evo.12281

Chamberlain, S.A., Bronstein, J.L., Rudgers, J.A., 2014. How context dependent are species interactions? Ecol Lett 17, 881–890. https://doi.org/10.1111/ele.12279

Chesson, P., 2000. Mechanisms of maintenance of species diversity. Ann. Rev. Ecol. Syst. https://doi.org/10.2307/221736

Chmielewski, M.W., Khatchikian, C., Livdahl, T., 2010. Estimating the per Capita Rate of Population Change: How Well Do Life-History Surrogates Perform? Annals of the Entomological Society of America 103, 734–741. https://doi.org/10.1603/AN09162

Costanzo, K.S., Westby, K.M., Medley, K.A., 2018. Genetic and environmental influences on the size-fecundity relationship in Aedes albopictus (Diptera: Culicidae): Impacts on population growth estimates? PLoS ONE 13, e0201465. https://doi.org/10.1371/journal.pone.0201465

Couret, J., Dotson, E., Benedict, M.Q., 2014. Temperature, Larval Diet, and Density Effects on Development Rate and Survival of Aedes aegypti (Diptera: Culicidae). PLOS ONE 9, e87468. https://doi.org/10.1371/journal.pone.0087468

Fey, S.B., Cottingham, K.L., 2011. Linking biotic interactions and climate change to the success of exotic Daphnia lumholtzi. Freshwater Biology 56, 2196–2209. https://doi.org/10.1111/j.1365-2427.2011.02646.x

Gillooly, J.F., Charnov, E.L., West, G.B., Savage, V.M., Brown, J.H., 2002. Effects of size and temperature on developmental time. Nature 417, 70–73. https://doi.org/10.1038/417070a

Gilotra, S.K., Rozeboom, L.E., Bhattacharya, N.C., 1967. Observations on possible competitive displacement between populations of Aedes aegypti Linnaeus and Aedes albopictus Skuse in Calcutta. Bull World Health Organ 37, 437–446.

Golding, N., Nunn, M.A., Purse, B.V., 2015. Identifying biotic interactions which drive the spatial distribution of a mosquito community. Parasites & Vectors 8, 367. https://doi.org/10.1186/s13071-015-0915-1

Grainger, T.N., Rego, A.I., Gilbert, B., 2018. Temperature-Dependent Species Interactions Shape Priority Effects and the Persistence of Unequal Competitors. The American Naturalist 191, 197–209. https://doi.org/10.1086/695688

Hart, S.P., Freckleton, R.P., Levine, J.M., 2018. How to quantify competitive ability. Journal of Ecology 106, 1902–1909. https://doi.org/10.1111/1365-2745.12954

Hartig, F., 2019. DHARMa: Residual Diagnostics for Hierarchical (Multi-Level / Mixed) Regression Models.

Hastings, A., 1980. Disturbance, coexistence, history, and competition for space. Theoretical Population Biology.

Inouye, B.D., 2001. Response Surface Experimental Designs for Investigating Interspecific Competition. Ecology 82, 2696–2706. https://doi.org/10.2307/2679954

Joe, H., Zhu, R., 2005. Generalized Poisson Distribution: the Property of Mixture of Poisson and Comparison with Negative Binomial Distribution. Biometrical Journal 47, 219–229. https://doi.org/10.1002/bimj.200410102

Juliano, S., O’Meara, G., Morrill, J., Cutwa, M., 2002. Desiccation and thermal tolerance of eggs and the coexistence of competing mosquitoes. Oecologia 130, 458–469. https://doi.org/10.1007/s004420100811

Juliano, S.A., 2010. Coexistence, Exclusion, or Neutrality? A Meta-Analysis of Competition between Aedes Albopictus and Resident Mosquitoes. Israel Journal of Ecology & Evolution 56, 325–351. https://doi.org/10.1560/IJEE.55.3-4.325

Juliano, S.A., 2009. Species Interactions Among Larval Mosquitoes: Context Dependence Across Habitat Gradients. Annu. Rev. Entomol. 54, 37–56. https://doi.org/10.1146/annurev.ento.54.110807.090611

Juliano, S.A., Lounibos, L.P., 2005. Ecology of invasive mosquitoes: effects on resident species and on human health. Ecol Lett 8, 558–574. https://doi.org/10.1111/j.1461-0248.2005.00755

Juliano, S.A., Ribeiro, G.S., Maciel-de-Freitas, R., Castro, M.G., Codeço, C., Lourenço-de-Oliveira, R., Lounibos, L.P., Juliano, S.A., Ribeiro, G.S., Maciel-de-Freitas, R., Castro, M.G., Codeço, C., Lourenço-de-Oliveira, R., Lounibos, L.P., 2014. She’s a femme fatale: low-density larval development produces good disease vectors. Memórias do Instituto Oswaldo Cruz 109, 1070–1077. https://doi.org/10.1590/0074-02760140455

Kaufman, M.G., Fonseca, D.M., 2014. Invasion Biology of Aedes japonicus japonicus(Diptera: Culicidae). Annu. Rev. Entomol. 59, 31–49. https://doi.org/10.1146/annurev-ento-011613-162012

Kesavaraju, B., Afify, A., Gaugler, R., 2012. Strain Specific Differences in Intraspecific Competition in Aedes albopictus (Diptera: Culicidae). J Med Entomol 49, 988–992. https://doi.org/10.1603/ME11245

Kingsolver, J.G., Huey, R.B., 2008. Size, temperature, and fitness: three rules. Evolutionary Ecology Research 10, 251–268.

Koenraadt, C.J., Kormaksson, M., Harrington, L.C., 2010. Effects of inbreeding and genetic modification on Aedes aegypti larval competition and adult energy reserves. Parasites & Vectors 3, 92. https://doi.org/10.1186/1756-3305-3-92

Leisnham, P.T., Juliano, S.A., 2010. Interpopulation differences in competitive effect and response of the mosquito Aedes aegypti and resistance to invasion by a superior competitor. Oecologia 164, 221–230. https://doi.org/10.1007/s00442-010-1624-2

Livdahl, T.P., Sugihara, G., 1984. Non-linear interactions of populations and the importance of estimating per capita rates of change. The Journal of Animal Ecology 53, 573–580. https://doi.org/10.2307/4535

Lounibos, L.P., 2007. Competitive displacement and reduction. Journal Of The American Mosquito Control Association 23, 276–282.

Lounibos, L.P., 2002. Invasions by insect vectors of human disease. Annu. Rev. Entomol. 47, 233–266.

Lounibos, L.P., Bargielowski, I., Carrasquilla, M.C., Nishimura, N., 2016. Coexistence of Aedes aegypti and Aedes albopictus (Diptera: Culicidae) in Peninsular Florida Two Decades After Competitive Displacements. J Med Entomol 53, 1385–1390. https://doi.org/10.1093/jme/tjw122

Lounibos, L.P., Juliano, S.A., 2018. Where vectors collide: the importance of mechanisms shaping the realized niche for modeling ranges of invasive Aedes mosquitoes. Biol Invasions 20, 1913–1929. https://doi.org/10.1007/s10530-018-1674-7

Marcantonio, M., Metz, M., Baldacchino, F., Arnoldi, D., Montarsi, F., Capelli, G., Carlin, S., Neteler, M., Rizzoli, A., 2016. First assessment of potential distribution and dispersal capacity of the emerging invasive mosquito Aedes koreicus in Northeast Italy. Parasites & Vectors 1–19. https://doi.org/10.1186/s13071-016-1340-9

Mariappan, T., Thenmozhi, V., Udayakumar, P., Bhavaniumadevi, V., 2015. An observation on breeding behaviour of three different vector species (Aedes aegypti Linnaeus 1762, Anopheles stephensi Liston 1901 and Culex quinquefasciatus Say 1823) in wells in the coastal region of Ramanathapuram district, Tamil Nadu, India. International Journal of Mosquito Research 2, 42–44.

Medlock, J.M., Hansford, K.M., Schaffner, F., Versteirt, V., Hendrickx, G., Zeller, H., Bortel, W.V., 2012. A Review of the Invasive Mosquitoes in Europe: Ecology, Public Health Risks, and Control Options. Vector Borne Zoonotic Dis 12, 435–447. https://doi.org/10.1089/vbz.2011.0814

Miazgowicz, K.L., Mordecai, E.A., Ryan, S.J., Hall, R.J., Owen, J., Adanlawo, T., Balaji, K., Murdock, C.C., 2019. Mosquito species and age influence thermal performance of traits relevant to malaria transmission. bioRxiv 769604. https://doi.org/10.1101/769604

Monnerat, A.T., Machado, M.P., Vale, B.S., Soares, M.J., Lima, J.B.P., Lenzi, H.L., Valle, D., 2002. Anopheles albitarsis Embryogenesis: Morphological Identification of Major Events. Memórias do Instituto Oswaldo Cruz 97, 589–596. https://doi.org/10.1590/S0074-02762002000400026

Mordecai, E.A., Cohen, J.M., Evans, M.V., Gudapati, P., Johnson, L.R., Lippi, C.A., Miazgowicz, K., Murdock, C.C., Rohr, J.R., Ryan, S.J., Savage, V., Shocket, M.S., Ibarra, A.S., Thomas, M.B., Weikel, D.P., 2017. Detecting the impact of temperature on transmission of Zika, dengue, and chikungunya using mechanistic models. PLOS Neglected Tropical Diseases 11, e0005568. https://doi.org/10.1371/journal.pntd.0005568

Mordecai, E.A., Paaijmans, K.P., Johnson, L.R., Balzer, C., Ben-Horin, T., de Moor, E., McNally, A., Pawar, S., Ryan, S.J., Smith, T.C., Lafferty, K.D., 2013. Optimal temperature for malaria transmission is dramatically lower than previously predicted. Ecology Letters 16, 22–30. https://doi.org/10.1111/ele.12015

Murrell, E.G., Juliano, S.A., 2008. Detritus Type Alters the Outcome of Interspecific Competition Between *Aedes aegypti* and *Aedes albopictus* (Diptera: Culicidae). Journal of Medical Entomology 45, 375–383. https://doi.org/10.1603/0022-2585(2008)45[375:DTATOO]2.0.CO;2.

Park, T., 1954. Experimental Studies of Interspecies Competition .2. Temperature, Humidity, and Competition in 2 Species of Tribolium. Physiological Zoology 27, 177–238.

Parton, W.J., Logan, J.A., 1981. A model for diurnal variation in soil and air temperature. Agricultural Meteorology 23, 205–216. https://doi.org/10.1016/0002-1571(81)90105-9

R Core Team, 2018. R: A language and environment for statistical computing. R Foundation for Statistical Computing, Vienna, Austria.

Raminani, L.N., Cupp, E.W., 1978. Embryology of Aedes aegypti (L.) (Diptera: Culicidae): Organogenesis. International Journal of Insect Morphology and Embryology 7, 273–296. https://doi.org/10.1016/0020-7322(78)90009-0

Reiskind, M.H., Zarrabi, A.A., 2012. Is bigger really bigger? Differential responses to temperature in measures of body size of the mosquito, Aedes albopictus. Journal of Insect Physiology 58, 911–917. https://doi.org/10.1016/j.jinsphys.2012.04.006

Schaffner, F., Kaufmann, C., Hegglin, D., Mathis, A., 2009. The invasive mosquito Aedes japonicus in Central Europe. Medical and Veterinary Entomology 23, 448–451. https://doi.org/10.1111/j.1365-2915.2009.00825.x

Seyfarth, M., Khaireh, B.A., Abdi, A.A., Bouh, S.M., Faulde, M.K., 2019. Five years following first detection of Anopheles stephensi (Diptera: Culicidae) in Djibouti, Horn of Africa: populations established&#8212;malaria emerging. Parasitol Res 118, 725–732. https://doi.org/10.1007/s00436-019-06213-0

Singh, P., Lingala, M.A.L., Sarkar, S., Dhiman, R.C., 2017. Mapping of Malaria Vectors at District Level in India: Changing Scenario and Identified Gaps. Vector-Borne and Zoonotic Diseases 17, 91–98. https://doi.org/10.1089/vbz.2016.2018

Sprenger, D., Wuithiranyagool, T., 1986. The Discovery and Distribution of Aedes albopictus in Harris County, Texas. JAMCA 2, 217–219.

Strayer, D.L., Eviner, V.T., Jeschke, J.M., Pace, M.L., 2006. Understanding the long-term effects of species invasions. Trends in Ecology & Evolution 21, 645–651. https://doi.org/10.1016/j.tree.2006.07.007

Surendran, S.N., Sivabalakrishnan, K., Sivasingham, A., Jayadas, T.T.P., Karvannan, K., Santhirasegaram, S., Gajapathy, K., Senthilnanthanan, M., Karunaratne, S.P., Ramasamy, R., 2019. Anthropogenic Factors Driving Recent Range Expansion of the Malaria Vector Anopheles stephensi. Front Public Health 7. https://doi.org/10.3389/fpubh.2019.00053

Takken, W., Lindsay, S., 2019. Increased Threat of Urban Malaria from Anopheles stephensi Mosquitoes, Africa - Volume 25, Number 7—July 2019 - Emerging Infectious Diseases journal - CDC. Emerging Infectious Diseases 25, 1431–1433. https://doi.org/10.3201/eid2507.190301

Thomas, S., Ravishankaran, S., Justin, J.A., Asokan, A., Mathai, M.T., Valecha, N., Thomas, M.B., Eapen, A., 2016. Overhead tank is the potential breeding habitat of Anopheles stephensi in an urban transmission setting of Chennai, India. Malaria Journal 15. https://doi.org/10.1186/s12936-016-1321-7

Verdonschot, P.F.M., Besse-Lototskaya, A.A., 2014. Flight distance of mosquitoes (Culicidae): A metadata analysis to support the management of barrier zones around rewetted and newly constructed wetlands. Limnologica 45, 69–79. https://doi.org/10.1016/j.limno.2013.11.002

Westby, K.M., Sweetman, B.M., Horn, T.R.V., Biro, E.G., Medley, K.A., 2019. Invasive species reduces parasite prevalence and neutralizes negative environmental effects on parasitism in a native mosquito. Journal of Animal Ecology 88, 1215–1225. https://doi.org/10.1111/1365-2656.13004

Wilke, A.B.B., Benelli, G., Beier, J.C., 2020. Beyond frontiers: On invasive alien mosquito species in America and Europe. PLOS Neglected Tropical Diseases 14, e0007864. https://doi.org/10.1371/journal.pntd.0007864

Yadav, R., Tyagi, V., Sharma, A.K., Tikar, S.N., Sukumaran, D., Veer, V., 2017. Overcrowding Effects on Larval Development of Four Mosquito Species Aedes Albopictus, Aedes Aegypti, Culex Quinquefasciatus and Anopheles Stephensi. International Journal of Research Studies in Zoology 3. https://doi.org/10.20431/2454-941X.0303001

Yee, D.A., Kaufman, M.G., Juliano, S.A., 2007. The significance of ratios of detritus types and micro-organism productivity to competitive interactions between aquatic insect detritivores. Journal of Animal Ecology 76, 1105–1115. https://doi.org/10.1111/j.1365-2656.2007.01297.x

